# Spatial transcriptomics reveals modulation of transcriptional networks across brain regions after auditory threat conditioning

**DOI:** 10.1101/2024.09.25.614979

**Authors:** Joy Otten, Shu Dan, Luise Rostin, Alex E. Profetto, Roy Lardenoije, Torsten Klengel

## Abstract

Prior research has demonstrated genome-wide transcriptional changes related to fear and anxiety across species, often focusing on individual brain regions or cell types. However, the extent of gene expression differences across brain regions and how these changes interact at the level of transcriptional connectivity remains unclear. To address this, we performed spatial transcriptomics RNAseq analyses in an auditory threat conditioning paradigm in mice. We generated a spatial transcriptomic atlas of a coronal mouse brain section covering cortical and subcortical regions, corresponding to histologically defined regions. Our finding revealed widespread transcriptional responses across all brain regions examined, particularly in the medial and lateral habenula, and the choroid plexus. Network analyses highlighted altered transcriptional connectivity between cortical and subcortical regions, emphasizing the role of steroidogenic factor 1. These results provide new insights into the transcriptional networks involved in auditory threat conditioning, enhancing our understanding of molecular and neural mechanisms underlying fear and anxiety disorders.

## Introduction

Stress- and anxiety disorders lack defining macroscopic or microscopic brain pathologies. However, previous work suggested changes at the systems biology and neural circuit level associated with fear, stress, and anxiety(*1*). Notably, genetic, epigenetic, and transcriptomic studies have highlighted a dysregulation in glucocorticoid signaling, inflammatory response, and metabolic processes related to stress and anxiety(*2*). Recent progress in understanding the underlying neurocircuitry associated with stress- and anxiety disorders in humans and animal models have shown that multiple brain regions contribute to the molecular, physiological, and behavioral effects of stress and anxiety(*1, 3, 4*). While the amygdala and its subnuclei are key nodes of the fear neurocircuitry(*5–7*), a much wider set of brain regions participate in this circuit, including the hippocampus(*6, 8–10*), medial prefrontal cortex(*11–14*), thalamus(*15, 16*), and brain stem regions(*17*), all contributing to the pattern of behavioral and physiological consequences of stress and anxiety.

Evidence suggests that maladaptive transcriptional changes within relevant brain regions are central to many stress- and anxiety related disorders(*18–20*). However, the contribution of system-wide gene expression differences across brain regions to behavioral outcomes remains underexplored. We use auditory threat conditioning (aTC), a robust behavioral paradigm to investigate aspects of learning, memory, and stress responses, to better understand the coordinated transcriptomic changes across multiple brain regions *in situ* using spatial transcriptomics. *In situ* spatial RNAseq technologies enable the profiling of genome-wide gene expression within the morphological context of tissue sections, overcoming prior limitations of bulk- and single-cell RNAseq technologies(*21*). Our results emphasize a non-overlapping transcriptional response of each brain region to aTC, highlight brain regions with particularly strong gene expression changes, and uncovered networks of transcriptional connectivity revealing a deeper insight into brain networks associated with threat conditioning.

## Results

### Unsupervised clustering of stRNAseq data resolves brain region anatomy

To investigate brain region- and group-specific transcriptional profiles after aTC, we generated a molecular atlas of a coronal section of the mouse brain targeting the hippocampus, amygdala, and thalamic regions using 10X Visium Spatial Transcriptomics RNA sequencing (stRNAseq) (**Figure 1A**). Across all samples (n=16), we identified n=46,837 spatially barcoded spots carrying whole-genome gene expression information, detecting n=19,428 unique genes with an average of n=4,041 genes per spot after filtering (**Supplemental Figure 1A-B**). We established an unsupervised clustering method to create an anatomical map of brain subregions based on stRNAseq data. This involved applying principal component analysis (PCA) for dimensionality reduction and Leiden clustering and led to the identification of 35 distinct brain regions (**Figure 1B-C**). The two-step clustering process first established 11 clusters, corresponding to broad brain regions, including the hippocampal formation, thalamus, and cortical regions, congruent with the known anatomy of the mouse brain (**Supplemental Figure 1C-D**). Subsequent re-clustering of the broader-defined regions led to the identification of 35 subclusters, corresponding to cortical layers (L1, L2/3, L4, L5, and L6), as well hippocampal subfields including CA1/2, CA3, the dentate gyrus, among others (**Figure 1B-C, Supplemental Figure 1E-F**). Unsupervised clustering based on the underlying stRNAseq data detected technical variations introduced presumably due to small discrepancies in the anterior-posterior axis or slight tilting during sectioning of the brains (**Supplemental Figure 2**). The resulting variability in the data led to the formation of two clusters only represented in a subset of samples, which were discarded from downstream analyses, resulting in 33 clusters representing distinct brain regions across all samples.

**Figure 1.**
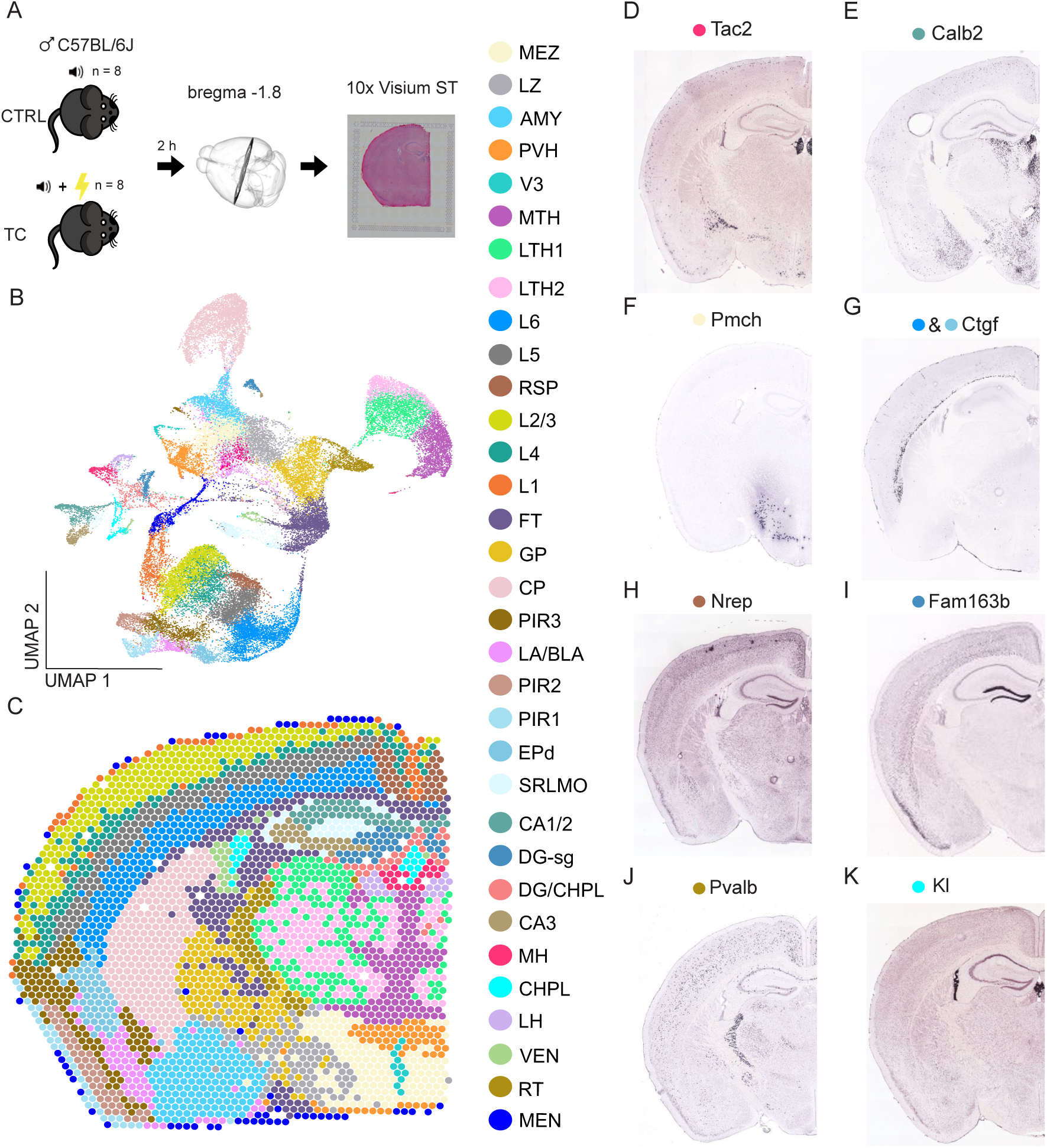
Unsupervised clustering of stRNAseq data. (**A**) Schematic representation of aTC experiment (total n=16 male C57BL/6J mice) with n=8 control and n=8 mice exposed to aTC. Mice were sacrificed 2 hours after exposure, and coronal brain sections at approximately bregma -1.8 were collected. The left hemisphere was placed on a 10X Visium chip and processed. (**B**) Uniform Manifold Approximation and Projection for Dimension Reduction (UMAP) plot illustrating the highest variance in the data based on principal components, effectively representing the separation of individual brain regions based on stRNAseq data from all samples. Colors represent different brain regions with color codes on the right. (**C**) Representative example of 2D reconstruction of stRNAseq data showing 33 distinct clusters. (**D-K**) Validation stRNAseq clustering with region-specific markers overlapping with representative markers derived from the Allen Brain Atlas (ABA). Abbreviations for brain regions derived from ABA. MEZ: Hypothalamic medial zone, LZ: Hypothalamic lateral zone, AMY: Amygdala, PVH: Paraventricular hypothalamic nucleus, V3: third ventricle, MTH: Medial thalamus, LTH1: Lateral thalamus 1, LTH2: Lateral thalamus 2, L6: Cortical layer 6, L5: Cortical layer 5, RSP: Retrosplenial cortex, L2/3: Cortical layer L2/3, L4: Cortical layer 4, L1: Cortical layer 1, FT: Fiber tract, GP: Globus pallidus, CP: Caudoputamen, PIR3: Piriform area polymorph layer 3, LA/BLA: Lateral amygdalar nucleus and basolateral amygdalar nucleus, PIR2: Piriform area polymorph layer 2, PIR1: Piriform area polymorph layer 1, EPd: Endopiriform nucleus, SRLMO: includes stratum radiatum, stratum lacunosum-moleculare, stratum oriens and stratum lucidum, CA1/2: Hippocampal region CA1 and CA2, DGsg: Dentate gyrus granule cell layer, DG/CHPL: Dentate gyrus and choroid plexus, CA3: Hippocampal region CA3, MH: Medial habenula, CHPL: Choroid plexus, LH: Lateral habenula, VEN: Ventricle, RT: Reticular nucleus of the thalamus, MEN: Meninges.

To validate the clustering approach, a differential expression analysis between brain regions was performed, identifying brain region-specific marker genes (Wilcoxon ranked sum test, p_FDR_<0.05, n=16 per brain region) (**Supplemental Figure 3, Supplemental Table 1**). *In situ* hybridization data from the Allen Brain Atlas (ABA) showed strong overlap with marker genes associated with regions across cortical and subcortical areas(*22*). For example, we validate the expression of *Tac2* and *Calb2*, the top marker genes for the medial and lateral habenula, respectively (**Figure 1D-K, Supplemental Figure 3)**. Additional regions validated by comparison with the ABA include the hypothalamic medial zone (*Pmch*), cortex layer 6 (*Ctgf*), retrosplenial cortex (*Nrep*), endopiriform cortex (*Ctgf*), dentate gyrus, granule cell layer (*Fam163b*), the reticular nucleus of the thalamus (*Pvalb*), and the choroid plexus (*Kl*). In summary, unsupervised clustering of stRNAseq data defined brain regions that correspond to known histologically and molecularly defined structures.

### Widespread transcriptional response to threat conditioning identified by stRNAseq: Beyond individual regions and genes

Next, we used the stRNAseq data for comparative analyses between mice exposed to aTC and tone-only controls to investigate associated gene expression changes across all brain regions. Auditory threat conditioning resulted in significantly increased freezing behavior in the aTC group compared to the control group, which was exposed to the same paradigm except for the co-terminating electrical foot shock (repeated measures two-way ANOVA, tone x group interaction: F_4, 56_=33.30, p<0.05; Sidak’s multiple comparisons posthoc test, p<0.05; n=8 per group) (**Figure 2A**). An advantage of stRNAseq in the context of this study is the parallel profiling of gene expression *in situ* covering multiple brain regions simultaneously. Prior genome-wide studies were limited to one or a few regions at a time, typically by micro-dissecting regions of interest in combination with bulkRNAseq or snRNAseq. We applied a pseudo-bulk approach with a linear mixed model to identify differentially expressed genes (DEGs) between groups in all 33 brain regions in parallel. The analysis resulted in n=415 DEGs across all brain regions (**Figure 2B**, **Supplemental Figures 4-5, Supplemental Table 2,** p_FDR_<0.05, n=8 per group, independent of cell type composition) with the largest total number of DEGs in cortical layer L2/3. To account for differences in the number of spots per region, and thus artifacts of relative sequencing depth, we normalized the absolute number of DEGs by the number of spots per region (**Figure 2C**). The top 3 regions identified after accounting for the size of each region include the medial and lateral habenula and the choroid plexus, all previously implicated in threat conditioning or stress- and anxiety related disorders. DEGs found in the medial habenula include *Hspa5*, the circadian rhythm genes *Clock and Usp2*, as well as *Atg12*, an autophagy marker linked to stress responses (**Figure 2D**). DEGs in the lateral habenula included *Impa1* encoding an inositol monophosphatase involved in phospholipase C signaling and *Adam10*, a metallopeptidase involved in synaptogenesis and plasticity (**Figure 2E**). In the choroid plexus, DEGs included *Ncl*, involved in angiogenesis and endothelial function, *Kras*, part of the RAS/MAPK pathway, *Lama5*, a member of the laminin family regulating neurodevelopmental processes, and *Pdlim2*, which is involved in cell adhesion and migration (**Figure 2F, Supplemental Table 2**).

**Figure 2.**
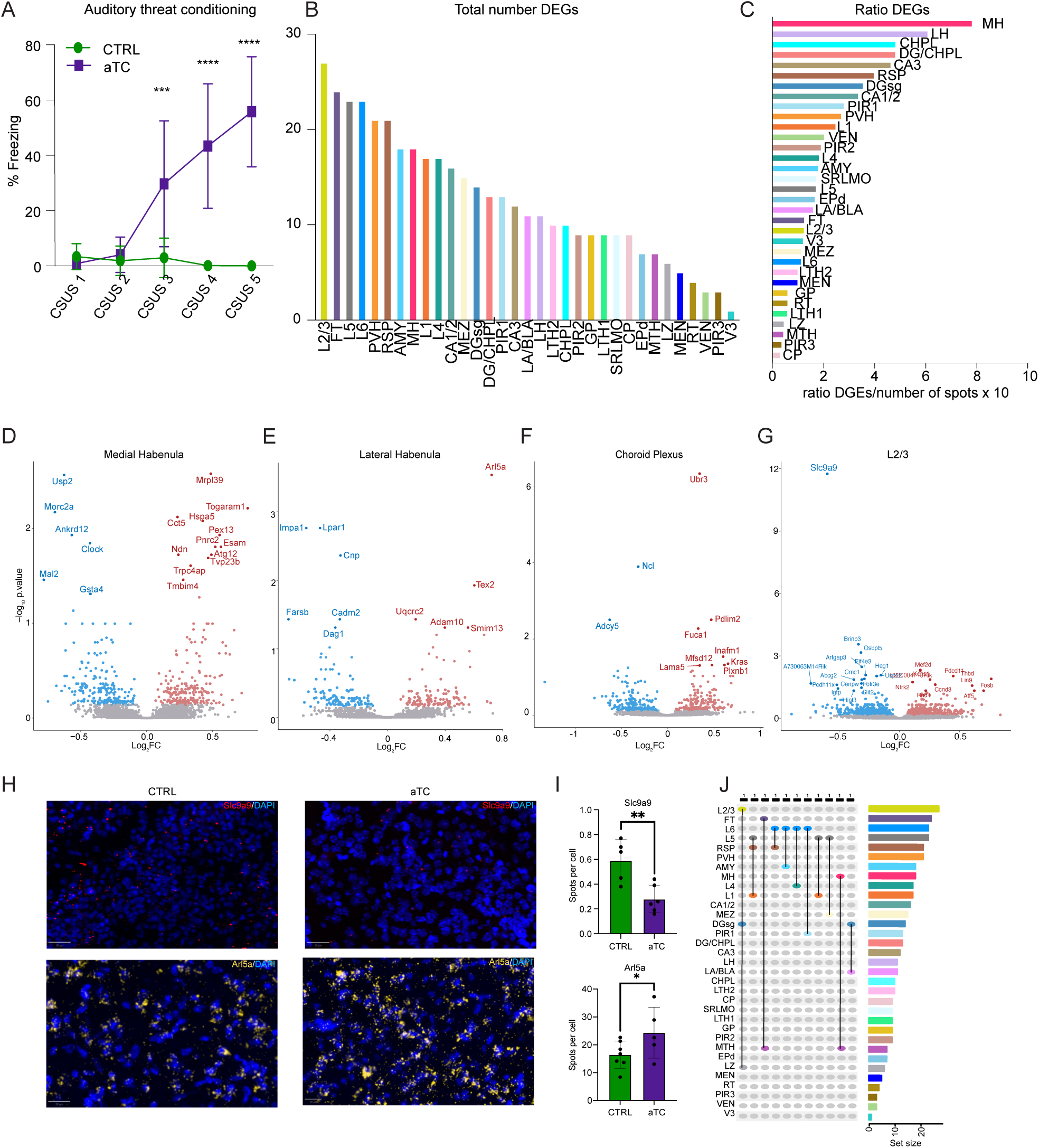
aTC engages most brain regions, inducing unique genome-wide transcriptional differences. (**A**) aTC data displaying the 5 tone-shock pairings (conditioned stimulus – unconditioned stimulus (CSUS) 1-5, x-axis) and percentage of time spent freezing on the y-axis. aTC results in a significant increase in freezing % in the threat-conditioned group after the second tone-shock pairing compared to the tone-control group. The data plotted is mean + SD, *** = p<0.0005; **** = p<0.0001. (**B**) 415 differentially expressed genes (DEGs) across 33 brain regions (p_FDR_ < 0.05), with L2/3, FT, and L5 with the largest number of total DEGs. Pseudo-bulk limma model (group + slide) per brain region. (**C**) DEGs normalized to the number of spots per region suggest that the medial and lateral habenula as well as the choroid plexus as top 3 most responsive brain regions. (**D-G**) Representative volcano plots showing DEGs. on the x-axis the log_2_ fold change is presented, and on the y-axis the -log_10_ p.value. DEGs in the medial habenula (**D**), lateral habenula (**E**), choroid plexus (**F**), and cortical layer 2/3 (**G**). Upregulated genes in red; downregulated genes in blue. Colored dots with gene names represent DEGs with p_FDR_<0.05. The remaining-colored dots represent nominal significant DEGs at p_nominal_<0.05. (**H**) Representative RNAscope images of *Slc9a9* (L2/3) or *Arl5a* (MH) expression after aTC. Scale bar represent 50µm (*Slc9a9*) or 20µm (*Arl5a*), respectively. (**I**) Quantification of RNAscope images of *Slc9a9* (n=6 per group) and *Arl5a* (n=7 for control and n=5 for aTC). ** = p<0.01; * = p<0.05. (**J**) Upset plot showing minimal overlap of DEGs between brain regions. On top the number of DEGs that overlap between brain regions is indicated.

To validate the technical aspects of stRNAseq we selected the top DEG from L2/3 (*Slc9a9*) and the lateral habenula (*Arl5a*), respectively. RNAscope *in situ* hybridization replicated the downregulation of *Slc9a9* and the upregulation of *Arl5a* in an independent cohort of mice supporting the reproducibility of the stRNAseq data obtained (**Figure 2E, G-I, Supplemental Figure 6,** one-tailed t-test, p<0.05; analysis of freezing behavior in the replication cohort: repeated measures two-way ANOVA, tone x group interaction: F_4, 52_=23.14, p<0.05; Sidak’s multiple comparison posthoc test, p<0.05, n_ctrl_=7, n_aTC_=8).

Across all regions investigated, we detected immediate early genes (IEGs), such as *Homer1*, *Egr3*, *Egr1*, and *Fosb*, congruent with the sampling timepoint of 2 hours after aTC (p_FDR_<0.05, **Supplemental Table 2**). We also detect genes previously reported in the context of threat conditioning, fear, stress, anxiety, and memory formation including -among others-*CamK2G* in L5, *Pde8b* and *Yy1* in L4, *Fkbp4* and *Klf10* in the dentate gyrus, *Gad1* in the molecular layer of the piriform area, *Ntrk2* and *Eif4e3* in L2/3, *Camk2n1* in the retrosplenial cortex, *Gpr12* and *Fkbp5* in the amygdala, *Ppp1r1b* (DARPP-32) in the lateral ventricle/subependymal zone, and *Ctnnb1* in the globus pallidus.

Predictably, we also discovered a plethora of genes without prior evidence for a role in threat conditioning; however, some have been implicated in other aspects of emotion regulation or relevant molecular pathways. (**Figure 2B-C, Supplemental Table 2**). For example, *Brinp3* in L2/3 and *Brinp2* in the polymorph layer of the piriform area have been implicated in anxiety and hyperactivity, although their role in fear learning remains unclear. *Aldh5a1*, which encodes the enzyme succinate semialdehyde dehydrogenase (SSADH) catalyzing the conversion of GABA and GHB to succinate, which subsequently enters the tricarboxylic acid cycle and is upregulated in the dentate gyrus. We also detected a substantial number of DEGs in white matter tracts (encompassing the corpus callosum, fimbria, stria terminalis, and internal capsule) including *Plat*, a modulator of synaptic plasticity, and *Tmem163*, which is involved in P2X receptor signaling (**Supplemental Table 2**).

In general, DEGs were distributed across all brain regions, suggesting that aTC affects gene expression changes beyond previously investigated structures and across the entire brain section. Although we found many genes related to various aspects of brain function, fear, anxiety, and memory formation, each brain region exhibited a unique set of DEGs with minimal overlap with other brain regions (**Figure 2J, Supplemental Table 2**). A small fraction of all DEGs correlated with freezing (26/415 DEGs, p_Nominal_<0.05), the main phenotypic readout of aTC in this study, suggesting a direct role for these genes related to freezing behavior (**Supplemental Figure 7A-D, Supplemental Table 3**). For example, *Pgd* in the polymorph layer of the piriform area showed a negative correlation with freezing, while *Rnd1* in L2/3 and *Hagh* in the endopiriform cortex exhibited positive correlations.

To investigate global changes in transcriptional regulation, we performed gene set enrichment analyses (GSEA) on all nominal significant DEGs (p_Nominal_<0.05) per brain region (**Supplemental Table 4**). Similar to the individual DEGs, each brain region produced sets of enriched terms with minimal overlap, supporting the idea that aTC results in a unique, region-specific transcriptional response. In summary, our data suggest a brain-wide but region-specific transcriptional response to aTC, with unique genes and pathways regulated in each brain region.

### From individual brain regions and genes to transcription-based networks of threat conditioning

Prior work firmly established the involvement of a circuit including the amygdala, hippocampus, and medial prefrontal cortex in aTC. However, additional brain regions likely contribute to the behavioral response to aTC in a time-dependent fashion with different brain regions engaging at different time points during and after the paradigm(*23*). Our data support this, however, our knowledge of the interaction of brain regions at the gene expression level in aTC remains scarce. This contrasts with functional imaging analyses using magnetic resonance imaging (MRI), which can infer dynamic response of brain activity and identify patterns of co-activation, known as functional connectivity. While whole-brain functional connectivity analyses via MRI are common in humans, this remains challenging in freely moving rodent models. However, recent studies using whole-brain immediate early gene expression (IEGs, such as *c-Fos* and *Arc*), which track neuronal activity, have helped to determine functional coactivation patterns at the single gene expression level, adding complimentary information to functional- and structural connectivity studies(*24–26*).

To gain insight into functional connectivity pattern in aTC, we first performed weighted gene co-expression analysis (WGCNA) across all 33 brain regions simultaneously. To prevent the detection of inherent differences between brain regions, we regressed the effects of brain region identity out. WGCNA identified 28 gene modules, with 2 modules showing a significant positive correlation with group (CTRL vs. aTC.; p_FDR_<0.05, **Figure 3A-B**). The darkgrey module showed enrichment for the transcription factor E2F1, and the lightgreen module was enriched for the SF-1 (steroidogenic factor 1) transcription factor suggesting a regulation of transcriptional activity across brain regions by E2F1 and SF-1 (**Figure 3C-D, Supplemental Table 5**). The hub gene for the darkgrey module is *Bc1*, encoding a small non-messenger RNA, with prior evidence for a role in anxiety behavior. The hub gene for the lightgreen module is *Eef1a2* encoding for eukaryotic translation elongation factor 1 which is associated with neurodevelopmental disorders, epilepsy, and intellectual disability.

**Figure 3.**
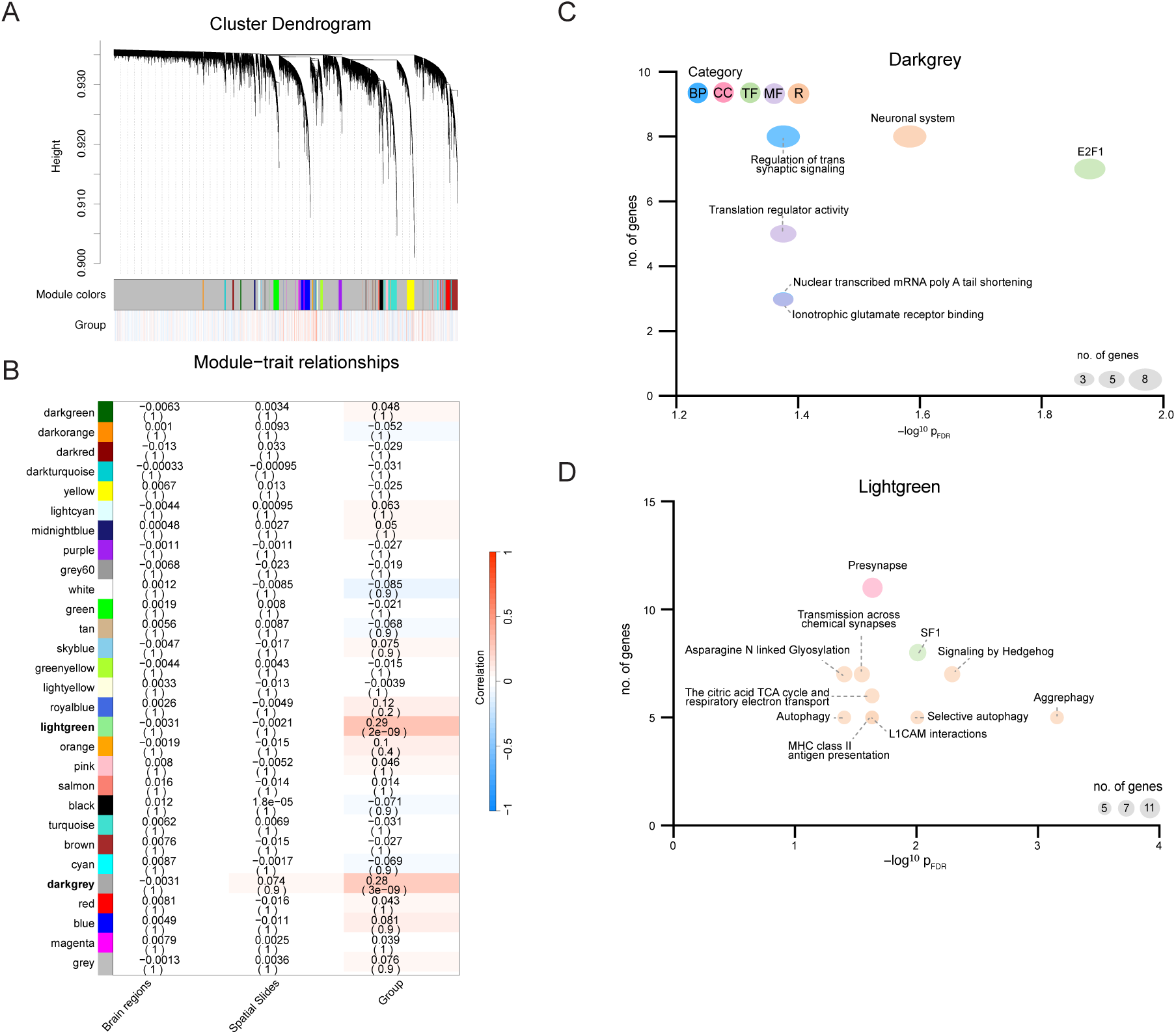
WGCNA suggests a role for E2F1 and SF-1. (**A**) Cluster dendrogram of a signed network revealing 28 modules. A large proportion of genes is not assigned to a specific module and thus cluster in the grey module. (**B**) Module-trait relationship showing 2 modules with a significant positive correlation with group (p_FDR_<0.05). No significant correlation to either brain region or Visium slides. (**C-D**) Bubble plots depicting selected enriched terms from the enrichment analysis in the darkgrey (**C**) or lightgreen (**D**) module, highlighting E2F1 and SF-1 enrichment in the darkgrey and lightgreen module, respectively. The x-axis represents -log^10^ FDR p-values and y-axis represents the number of genes contributing to the enriched term as well as the size of the bubble. The colors represent the different databases. BP: Gene ontology biological processes, CC: Gene ontology cellular component, TF: Transcription factor, MF: Gene ontology molecular functions, R: Reactome database.

To further explore the interaction of brain regions at the level of gene expression utilizing the unique capabilities of the stRNAseq dataset, we employed a strategy developed by Gandal et al., which utilizes expression differences between brain regions due to developmental patterning, differences in connectivity, and function, to define a unique transcriptomic regional identity(*27*). We adapted the method of Gandal et al. to identify transcriptional connectivity differences across all 33 brain regions. Seven pairs of cortical and subcortical regions (out of 528 pairs) exhibited significant attenuation in aTC compared to controls (**Figure 4A**, p_FDR_<0.05). At the gene expression level, attenuation indicates a reduction in gene expression differences between two brain regions in aTC compared to controls, suggesting that these two regions become transcriptionally more similar to each other after aTC. Significant attenuation was detected between the piriform area and globus pallidus, the reticular nucleus of the thalamus (RT), white matter fiber tracts, and the CA3 region (**Figure 4A**). Conversely, our analysis also shows over-patterning, indicating an increase in gene expression differences between two regions, suggesting that these regions become transcriptionally more dissimilar in aTC. Nine pairs exhibited significant over-patterning in aTC compared to controls. Over-patterning was detected between cortical layers L2/3, L4, L5, and L6, capturing the dorsal auditory and somatosensory areas as well as the barrel cortex and the hypothalamus. Furthermore, significant over-patterning was observed between L4 and the lateral ventricle, L4 and the dorsal endopiriform cortex (EPd), retrosplenial cortex and dentate gyrus, and the thalamus and dentate gyrus (**Figure 4B**, p_FDR_<0.05).

**Figure 4.**
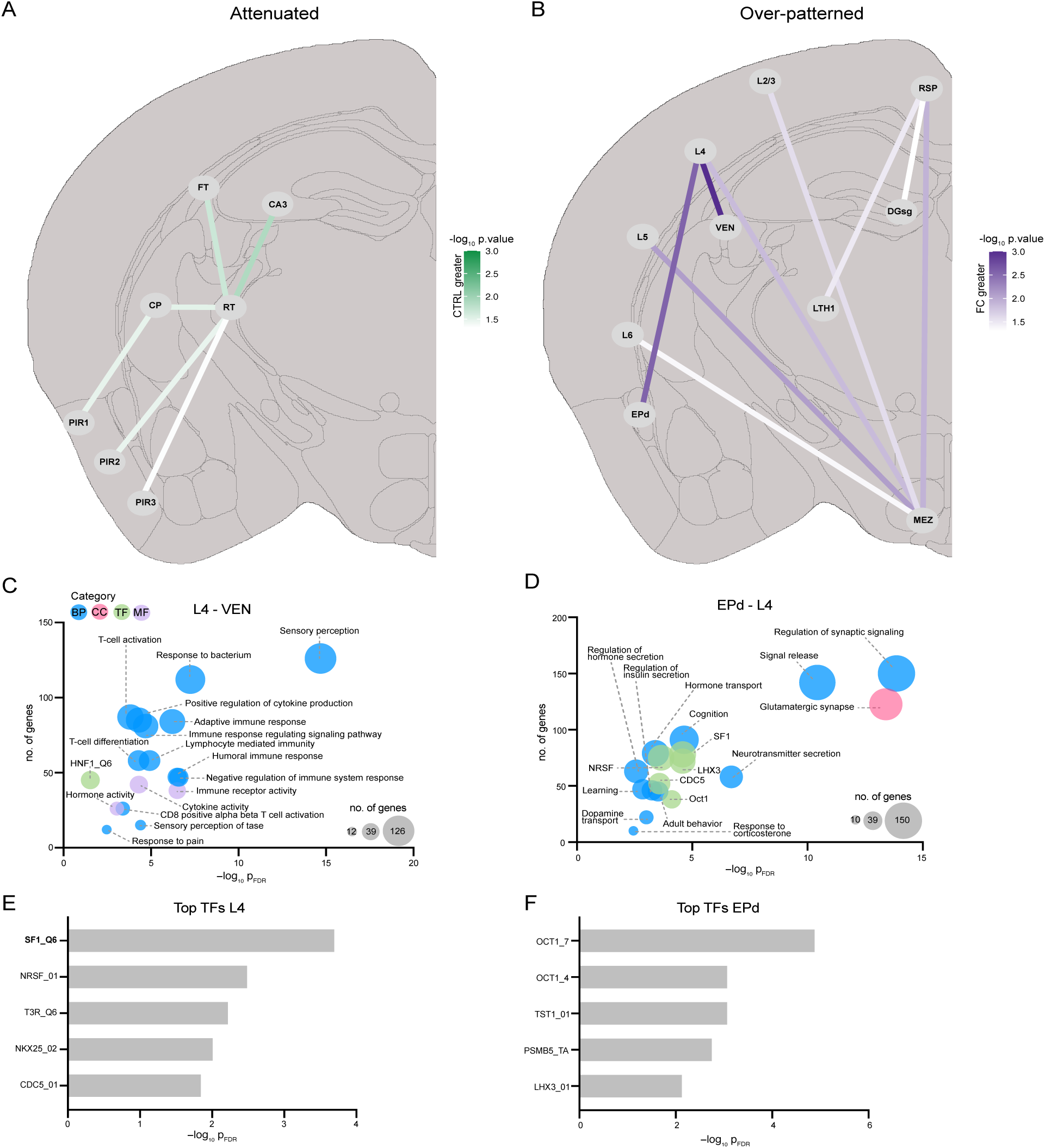
Transcriptional connectivity networks. (**A**) Attenuated network after aTC include region-to-region pairing between RT-PIR2, RT-PIR3, RT-CP, CP-PIR1, RT-FT (p=0.02), and RT-CA3 (p=0.013) with the latter two showing the strongest attenuation. (**B**) Over-patterned network after aTC include region-to-region pairing between RSP-MEZ, RPS-DGsg, RSP-LTH1, L2/3-MEZ, L6-MEZ, L5-MEZ, L4-VEN, and L4-EPd, with the latter two showing the strongest over patterning (p<0.001, p<0.01 respectively). (**C-D**) Bubble plots showing selected enriched terms of genes contributing to transcriptional connectivity changes between L4-VEN (**C**) or L4-EPd (**D**). Of note, differences between L4-EPd may be driven by SF-1. The x-axis represents -log^10^ FDR p-values and y-axis represents the number of genes contributing to the enriched term as well as the size of the bubble. The colors represent the different databases. BP: Gene ontology biological processes, CC: Gene ontology cellular component, TF: Transcription factor, MF: Gene ontology molecular functions, R: Reactome database. (**E-F**) Bar graph depicting the top 5 transcription factors of genes contributing to gene expression in L4 (**E**) or EPd (**F**). The x-axis represents the -log^10^ FDR and the y-axis depicts the transcription factors. SF-1 is specifically detected in L4.

To investigate the biological processes contributing to changes in transcriptional connectivity, we performed a pathway and transcription factor enrichment analysis on genes contributing to the attenuated or over-patterned network, respectively. Enrichment analysis yielded significant results for the attenuated connection between the RT and the polymorph layer 3 of the piriform cortex (PIR3) showing enrichment for SNARE interactions in vesicular transport. In addition, the connection between RT and the polymorph layer 2 of the piriform cortex (PIR2) showed enrichment for MAPK6/4 and IL-1 signaling among others (**Supplemental Table 6,** p_FDR_<0.05).

Vice versa, the over-patterned network connections between L4 and the ventricle showed enrichment for immune-related terms and the HNF1 transcription factor (**Figure 4C, Supplemental Table 6,** p_FDR_<0.05). Notably, we observed an enrichment of SF-1 within the over-patterned network between the EPd and L4, consistent with previous findings on the expression of *Nr5a1*, which encodes SF-1 in L4 interneurons (**Figure 4D, Supplemental Table 6,** p_FDR_<0.05)(*28, 29*). To identify the brain region contributing to the SF-1 signal, we classified genes based on their expression levels as belonging to either the EPd or L4 regions. Enrichment analysis of the genes assigned to L4 or EPd revealed significant enrichment of SF-1 and OCT-1 transcription factors, respectively (**Figure 4E-F, Supplemental Table 7,** p_FDR_<0.05). Collectively, our data suggest alterations in transcriptional connectivity between brain regions following aTC.

## Discussion

The results presented here provide evidence for brain-wide transcriptional changes in response to auditory threat conditioning beyond known brain structures and previously identified differentially expressed genes and networks. We identified 415 genome-wide significant DEGs across 33 brain regions *in situ*, significantly expanding the current understanding of the transcriptional response to aTC(*3, 4*). Network analyses reveal significant differences in transcription-based connectivity, involving both previously studied regions and those not typically associated with threat conditioning.

Spatial transcriptomic RNA sequencing detects gene expression in intact tissue sections(*21, 30*), preserving the anatomical context, which is a significant advantage over technologies that use homogenized tissue or single cells. In this study, we applied the 10X Genomics Visium spatial transcriptomics platform to an aTC paradigm in male C57BL/6J mice targeting brain regions including the hippocampal formation, the amygdala, and the cortex. Unsupervised clustering of the stRNAseq data resulted in a precise representation of known anatomical regions and robust overlap with gene expression patterns obtained from the Allen Mouse Brain Reference Atlas(*22*), demonstrating that spatial transcriptomics can faithfully identify anatomical regions based on their molecular characteristics.

Utilizing stRNAseq, we performed an unbiased differential gene expression analysis across 33 brain regions. Like prior investigations of whole-brain IEG expression after cued threat conditioning in mice, we find that training engages most brain regions(*19, 20, 25, 31*). However, our results significantly expand prior knowledge on the transcriptional response to aTC. Notably, we provide evidence for widespread transcriptional changes across all identified brain regions, with many DEGs not previously described in the context of aTC. While cortical regions showed a high number of DEGs overall, this finding is influenced by the proportionally higher number of sequencing reads allocated to these larger regions. When normalizing the number of DEGs to region size, the medial habenula (MH) displayed the highest number of DEGs per spot, paralleled by the largest number of GO terms significantly enriched by GSEA. Differentially regulated genes in the MH include – among others – *Clock* and *Usp2* (related to circadian rhythm), chaperone-related genes *Hspa5* and *Cct5*, autophagy marker *Atg12*, *Pex12* (implicated in mitochondrial dysfunction and oxidative stress), and *Pnrc2* (a coactivator of nuclear receptors including the estrogen and glucocorticoid receptor). Although the habenula has been studied for its role in motivational states and cognition, relatively little is known about the function of the MH in aTC(*32*). Our findings also highlight other regions with sparse evidence for their role in aTC, such as white matter tracts, the choroid plexus, and the endopiriform nucleus. Interestingly, only a fraction of DEGs directly correlated with freezing, the main behavioral phenotype in this experiment, indicating that the majority of DEGs contribute to other aspects of aTC beyond freezing.

While DEGs can illuminate specific molecular pathways implicated in aTC within distinct brain regions, our results indicate a more complex transcriptional regulation across numerous brain regions. Using WGCNA, we identified two gene expression modules positively correlated with aTC. Both highlight gene pathways relevant to aTC and importantly show an enrichment of genes targeted by the transcription factors E2F1 and SF-1. *Nr5a1*, encoding SF-1, is specifically expressed in the ventromedial hypothalamus and L4 interneurons(*29*). SF-1 has been linked to fear responses, anxiety, emotional states, and feeding behaviors(*33, 34*), (*35*) and prior evidence suggests that it controls the expression of cannabinoid receptor 1 (CB1), brain-derived neurotrophic factor (BDNF), corticotrophin-releasing hormone receptor 2 (Crhr2), and enzymes converting pregnenolone to cortisol and aldosterone(*36, 37*).

To further model brain network changes based on gene expression data, we adapted recently developed methods that infer the transcriptional connectivity of brain regions, similar to concepts used in imaging studies(*38–41*). This involved comparing the unique gene expression profiles of individual brain region pairs between control animals and those exposed to aTC.

On one side, we detect an attenuated transcriptional network with attenuation defined as a decrease in gene expression differences between two brain regions, making them more similar in regard to their transcriptional profiles. Attenuation was detected between the piriform area, caudo-putamen, CA3, white matter fiber tracts, and RT. The piriform area is critical for processing and coding olfactory information, an important sensory cue for mice regarding orientation, communication, and survival(*42*). The caudo-putamen, part of the dorsal striatum, has been implicated in rodent threat conditioning and fear learning in humans(*43*). While evidence for direct connections between the piriform area and caudo-putamen is elusive, imaging studies suggest relevant functional connectivity as part of olfactory signaling pathways(*44*). Similarly, there is no data supporting direct interactions between the piriform area and RT. The RT is a key structure for thalamic inhibition regulating thalamocortical interactions critical for sensory processing, attention, and cognition(*45*). However, robust evidence suggests a key role for RT in threat conditioning, and our data support transcriptional connectivity between the RT, dorsal striatum, CA3, and white matter fiber tracts.

On the other side we detected an over-pattered network defined by an increase in gene expression differences, rendering regions more dissimilar based on their gene expression level. We observed over-patterning in response to aTC in cortical layers (L2/3, L4, L5, and L6), the hypothalamus, lateral ventricle, endopiriform nucleus, retrosplenial cortex, dentate gyrus, and thalamus. Strong evidence exists for bidirectional anatomical and functional connections between cortical layers and the hypothalamus in the context of the cortico-hypothalamic pathway, which is critical for regulating sensory cues, hormone secretion, autonomic functions, and behavior, bypassing the amygdala(*46*). The retrosplenial cortex, as the most caudal part of the cingulate cortex, is interconnected with the hippocampal formation and thalamus and is important for memory formation and spatial navigation(*47*). The over-patterning of regional identity between the RSP, dentate gyrus, and thalamus supports a role for these pathways in fear memory acquisition and stress response. While first evidence for direct connections between L4 and the endopiriform nucleus emerges(*48*), the role of the endopiriform nucleus in fear conditioning is not well understood, although some data suggest that it can modulate amygdala function(*49*). Although we cannot assign specific valence to attenuation and over-patterning, the detected changes suggest functional differences due to aTC, providing a foundation for further investigation into the molecular mechanisms and underlying circuitry. Interestingly, global enrichment analyses using WGCNA and specifically across network connections suggest that the transcription factor SF-1, encoded by *Nr5a1*, may have a general regulatory role integrating stress response and learning. Although most of the literature point to the expression of *Nr5a1* in the ventromedial hypothalamus, recent evidence suggest also expression in L4 interneurons(*29*). In fact, our analyses show that the expression of *Nr5a1* in L4 is driving the transcriptional connection between L4 and the endopiriform cortex.

In summary, this study provides evidence for extensive brain-wide transcriptional changes in response to aTC. Significant differences in transcription-based functional connectivity were observed, including both well-studied and unexplored brain regions. These results highlight the complexity of the brain’s response to aTC and suggests new avenues for understanding the molecular mechanisms underlying this process.

The integration of spatial transcriptomics with behavioral paradigms offers a powerful approach to map brain-wide transcriptional responses. This approach preserves the anatomical context of gene expression, allowing for a more comprehensive understanding of how different brain regions contribute to behavioral responses. The identification of DEGs and the mapping of transcription-based functional connectivity patterns provide new insights into the brain’s molecular and functional architecture in response to threat.

## Material and Methods

### Animals

Adult male C57BL/6J mice aged 9 weeks were obtained from The Jackson Laboratory (Bar Harbor, ME). Mice were group-housed in a temperature-controlled vivarium with set-point maintained at 22° C (±1°), a relative humidity controlled at 40–50%, and a 12-hour light-dark cycle with light on at 7am and light off at 7pm. Mice had access to standard rodent chow and water *ad libitum*. All procedures used were approved by the McLean Institutional Animal Care and Use Committee (IACUC) and in compliance with National Institutes of Health Guide for the Care and Use of Laboratory Animals.

### Auditory threat conditioning and tissue processing

Standard auditory threat conditioning(*50, 51*) using a 5 tone-shock paradigm was performed. After a 10-day acclimation period, mice were habituated to the experimental setup for 10 min for 2 consecutive days. In total, we used 16 mice (8 control, 8 threat conditioned). Threat conditioning was performed using white-light illuminated, standard rodent modular test chambers (ENV-008-VP; Med Associates Inc., St. Albans, VT). The paradigm consisted of five trials of a tone conditioned stimulus (CS; 30s tone, 6 kHz, 75 dB), which co-terminated with a mild foot-shock (1s, 0.6 mA) as the unconditioned stimulus (US). The Pre-CS period lasted 180s and a stable 90s inter-trial interval (ITI) was used between each CS-US pairing. Control animals were exposed to the same paradigm but did not receive a foot shock. The duration of freezing was measured automatically using FreezeFrame5 (Actimetrics, Wilmette, IL). Mice were returned to their cages after the paradigm and sacrificed 2 hours later via decapitation under brief isoflurane anesthesia. Brains were extracted, frozen on dry ice, and stored in CryoELITE tissue vials (DWK Life Sciences) at −80°C. Brains were sectioned on cryostat at 10mm according to the recommendations of the 10x Visium protocol (CG000239 Rev A and CG000240 Rev A) targeting the coronal section at bregma -1.8. Left hemispheres were mounted on the 10x Visium slides.

### 10x Genomics Visium spatial transcriptomics

Optimal conditions for tissue permeabilization were determined using the Visium Spatial Tissue Optimization kit according to manufacturer’s instruction resulting in a permeabilization time of 18 min. Slide processing and RNA sequencing was performed based on the instructions of the manufacturer (10x Genomics, Pleasanton, CA) at the Biomedical Research Center core sequencing facility at the University of British Columbia, Vancouver, Canada. Sequencing was carried out on an Illumina NextSeq 500 platform with a sequencing depth of approximately 100 million read pairs per sample.

### Behavior quantification and Statistical analysis

The data present are shown as mean + standard deviation (SD). Behavior data were analyzed using GraphPad Prism version 10.0.0. We performed a two-way ANOVA followed by Šidák’s multiple comparison posthoc test. P-values of < 0.05 were considered statistically significant.

### Fluorescent in situ hybridization (RNAscope)

The RNAscope Multiplex Fluorescent Reagent Kit V2 (cat. no. 323100, Advanced Cell Diagnostics) was performed according to the manufacturer’s instructions. Frozen mice brains were sliced at 20μm in the cryostat and dried for one hour at -20°C before being stored at -80°C. To prevent over-digestion of the samples while performing the RNAscope protocol Multiplex Fluorescent Reagent Kit v2 protocol the incubation time of the Protease IV was optimized to 15min. Opal dyes 570 (cat. no. FP1488001KT, Akoya Bioscience), and 690 (FP1497001KT, Akoya Bioscience) in a 1:1500 dilution were used to mark Channel 2-3. The probes used were Mm-Arl5a-C3 (cat. no. 1229731-C3) and Mm-Slc9a9-C2 (cat. no. 1229741-C2), which were designed by Bio-Techne GmbH. The slides were counterstained with DAPI (provided in the previously described kit), mounted with ProLong Gold Antifade Reagent (cat. No. P36931, Thermo Fisher Scientific), and stored at 4°C until image acquisition.

### Imaging and quantification

Whole slide images of the mouse brain were scanned at 20x magnification on an Olympus VS200 slide scanner at identical settings per RNAscope batch. Images were aligned with the Allen Brain Atlas (V3p1) coronal with the Fiji (2.14.0/1.54f) plugin. Annotations from the Allen Brain Atlas were loaded in QuPath(*52*). Thresholds that were used to detect nuclei stained with DAPI using the Cell detection feature were a requested pixel size of 0.3µm, a background radius of 30µm, a median filter radius of 2µm, a sigma of 1.5µm, a minimum area of 13µm^2^, a maximum area of 400µm^2^, a threshold of 2000, and a cell expansion of 10. The following boxes are checked: use opening by reconstruction, split by shape, include cell nucleus, smooth boundaries, and make measurements. RNAscope probes were detected with subcellular detection with various thresholds set per probe. Detection thresholds for Slc9a9, batch1: 2000, and batch 2: 2500 and detection thresholds for Arl5a, batch1: 4000 and batch 2: 2500. The following boxes were checked: smooth before detection, split by shape, split by intensity, and include cluster. The spot and cluster parameters were set as follows: expected spot size = 2mm^2^, min spot size = 0.5mm^2^, and max spot size = 6mm^2^. For each annotated object the sum of the number of spots estimated is divided by the total cells. We performed a one-way t-test where p-values < 0.05 were considered statistically significant.

### Sequence data processing

FASTQ files were processed by Space Ranger (v.1.0.0, 10x Genomics) with steps including spatial de-barcoding, read-alignment to the *Mus musculus* genome mm10 (v.3.0.0) and barcode/UMI-generation. The reads are generated by spaceranger count. All downstream data analysis was performed using R.4.1.2 and Seurat (v.4.0.2)(*53*) unless stated otherwise.

### Quality control, filtering and normalization

All samples were merged with the merge function in Seurat. Spots with a total of less than 200 gene counts were filtered out. Genes with less than 10 counts in total or detected in less than 2 spots were excluded as well. Additional genes including *Ttr*, *Hbb-bs*, *Hbb-bt*, *Hbb-y*, *Hba-a2* and *Hba-a1* were removed from the analysis due to their discontinuous patterns(*54, 55*). The filtered data was subsequently normalized using SCTransform without returning only the variable genes to remove batch effects.

### Dimensionality reduction, clustering, and sub-clustering

To reduce the noise in the data, principal component analysis (PCA) was carried out using the PCA function. Top 18 principal components (PCs) cumulatively preserved more than 90% variation in the data and among the top 18 PCs, the variations between each consecutive PC were more than 0.1%. Therefore, top 18 PCs were used as input for the FindNeighbors function. FindClusters function with Leiden algorithm(*56*) and igraph method was then applied to cluster spots with similar gene expression profiles. Various resolutions ranging from 0.1 to 1.8 were tested. Using clustree (v.0.4.3)(*57*), 11 clusters from a resolution of 0.2 were determined to be most closely resembling the known anatomy of the mouse brain after visual inspection.

To further define sub-clusters(*58*) within each cluster, we repeated the above steps to determine the number of PCs and resolution for each of the 11 clusters. 17 PCs per cluster were used for sub-clustering while the resolution parameter varies per cluster (res = 0.3: cluster 1, 2, 8; res = 0.2: cluster 3, 4, 7, 9; res = 0.1: cluster 5; res = 0: cluster 6, 10, 11). In total 35 sub-clusters were identified, and 2 clusters were removed from further analysis because they were not detected across all samples. The resulting clusters were assigned brain region labels based on their location and cluster-specific gene expression (*see identification of cluster specific genes*) compared to the Allen Mouse Brain Atlas.

### Identification of cluster specific genes

To identify cluster specific marker genes, the FindAllMarkers function was used to perform Wilcoxon Rank Sum test on genes that are detected in a minimum fraction of 25% spots in either of the two populations, logFC threshold of 0.25 and p_Nominal_ < 0.01. All p-values were adjusted for multiple testing through the Benjamini-Hochberg (BH) procedure.

### Differential expression analysis

Filtered unnormalized counts were pseudo-bulked by cluster to create a matrix similar to standard bulk RNAseq analyses. Differential gene expression analysis was performed using limma (v.3.50.3)(*59*). Pseudo-bulked counts were filtered again with filterByExpr(*60*) to remove genes with 0 count consistently across multiple samples. Scaling factors were calculated using calcNormFactors(*60*) with weighted trimmed mean of M-values (TMM)(*61*) to normalize and enable transformation in voom(*62*). A linear mixed effect model was used to determine the differentially expressed genes between groups while controlling for slide batch effects. Bacon(*63*) (v.1.22.0) was used for p-value inflation adjustment, and the FDR method was used to correct for multiple testing, genes with adjusted p-values < 0.05 were considered statistically significant.

### Gene set enrichment analysis

Gene set enrichment analysis (GSEA) was carried out using msigdbr (v.7.5.1)(*64, 65*) together with the enricher function from clusterProfiler (v.4.2.2)(*66*). Nominally significant differentially expressed genes (DEGs) per cluster (p_Nominal_ < 0.05) were used as input and all genes in the data were used as the universe (n=19,428). Q value cutoff was set at 0.05 and the resulting terms were corrected for multiple testing using the BH method. GSEA analysis was performed against three databases including Gene Ontology (GO)(*67*) Kyoto Encyclopedia of Genes and Genomes (KEGG)(*68*), Reactome(*69*), and for transcriptional connectivity analysis the TFT: transcription factor targets was used additionally. Specifically, the C3 regulatory target gene set category was used with the transcription factor targets legacy and the GTRD subset.

### Correlation analysis with freezing percentages

To identify genes correlated with freezing behavior we performed a Pearson correlation analysis (gene expression versus % freezing) on the DEGs (n=415). Percentage freezing is denoted as percentage freezing obtained from the 5^th^ CSUS pairing. In total 26 genes are significantly correlated (p_Nominal_ < 0.05) with freezing behavior.

### Cell type deconvolution

Deconvolution of stRNAseq leads to an estimated proportion of cell types per spot. To estimate cell type proportions per spot, we used CARD (v1.0)(*70*) together with previously published single-nucleus RNA sequencing (snRNAseq) data from adult male mice of the same age and strain(*71*). In total, n=300,422 cells were used for deconvolution. We derived data from the following brain regions: frontal and posterior cortex, entopeduncular nucleus, globus pallidus, hippocampus, striatum, and thalamus. The following cell types were included: astrocytes, endothelial cells, fibroblasts, GABAergic neurons, glutamatergic neurons, microglia, mural cells, oligodendrocytes, and polydendrocytes. To investigate cell type differences between groups, we used a two-way ANOVA test using Prism (v.10.0.0).

### Weighted gene co-expression network analysis

To identify genes with highly similar gene expression profiles we performed a weighted gene co-expression network analysis (WGCNA)(*72, 73*). WGCNA (v.1.72-1) utilizes a scale-free network to find gene co-expression modules based on correlation. Pseudo-bulked regional gene-expression data was used as input for normalization. Scaling factors were calculated using calcNormFactors^45^ and normalization was performed with compute counts per million (CPM) with prior.count = 0.5. First, we regressed out the effect of brain region identity along with slide effect, and sequencing depth with the empiricalBayesLM function. The signed network was constructed with the soft-power threshold of 4 to ensure a correlation coefficient above 0.9 with the Pearson correlation. The minimal module size was set to 30, the maximum block size to 30.000, the deepsplit to 2, and mergeCutHeight to 0.25. Of note out of the total of n=19,421 genes, n=14,591 genes were discarded for network analysis because they were not assigned to a module. This might be due to that most of these genes relate to the brain regions. The modules are correlated to the group (CTRL vs. aTC), brain regions, and Visium slides to construct the module-trait relationships.

### Transcriptional connectivity analysis

To identify transcriptional connectivity networks, we utilized a permutation-based method first described by Gandal et al(*27*). Briefly, the pseudo-bulked regional gene-expression data normalized by calcNormFactos with CPM was used as input, similar to the input for WGCNA described above. Next, we calculated DEGs between all possible brain region pairs 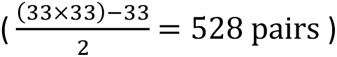 using a paired Wilcoxon rank-sum test in aTC and CTRL samples, separately. Genes with p_FDR_ < 0.05 were considered differentially expressed. The difference in the absolute number of DEGs per region between aTC and CTRL was defined as the ‘true difference’. Then, a permutation distribution of the absolute difference in DEGs between regions was generated to test the ‘true difference’. For each permutation (10,000 in total) ‘aTC’ and ‘CTRL’ status was randomly assigned to each sample with a consistent number of aTC and CTRL samples for each group. A two-tailed FDR corrected p-value was obtained from testing the ‘true difference’ against the permutation distribution. If the regional comparison yielded a p_FDR_ < 0.05 with the absolute number of DEGs between aTC < CTRL, then the regional comparison is considered significantly attenuated (both regions become transcriptionally more homogeneous after aTC). In turn, if the regional comparison yielded a p_FDR_ < 0.05 with the absolute number of DEGs between aTC > CTRL, the regional comparison was considered significantly over-patterned (both regions become transcriptionally more heterogeneous after aTC). To gain insight into the underlying molecular mechanisms, genes from the significant over-patterned or attenuated regional pairs were extracted. Filtering was applied to identify high-confidence genes from each significant attenuated or over-patterned pair identified by the permutation procedure mentioned above. Differentially expressed genes in each of the 10,000 permutations between each comparison were extracted. True differentially expressed genes between pairs of regions were calculated, ranging from a possibility of 0 to 10,000 occurrences. The true differentially expressed genes were present in less than 95% of their respective permutations. As outlined above aTC and CTRL labels were randomly assigned to samples during the permutation analysis.

With the assumption that the random assignment of the labels results in the detection of false positive genes, we took only DEGs that were present in less than 95% of the comparisons. Next, we performed enrichment and transcription factor enrichment analyses as described above to gain insights into the molecular mechanisms driven by DEGs between a pair of brain regions. Additionally, to investigate which brain region contributed to specific signals, we matched each gene to the brain region with higher expression levels. These gene sets were then used for an additional transcription factor enrichment analysis described above.

## Acknowledgments

The authors would like to thank Lakshmi Haferman for her assistance at the beginning of the project. JO was supported by the International Max Planck Research School for Genome Science. LR received support from the German Academic Scholarship Foundation.

## Author contributions

Conceptualization: JO, TK Methodology: JO, TK Investigation: JO, SD, LR, AP, RL, TK Visualization: JO Supervision: TK Writing: JO, TK

## Data and materials availability

Spatial transcriptomics data generated in this study is available at Gene Expression Omnibus (GEO) under accession number GSE243140. The analysis code is available from GitHub at https://github.com/klengellab/Spatialtranscriptomics_2024_aTC.

