## Supplemental Data for "Spatial transcriptomics reveals modulation of transcriptional networks across brain regions after auditory threat conditioning"

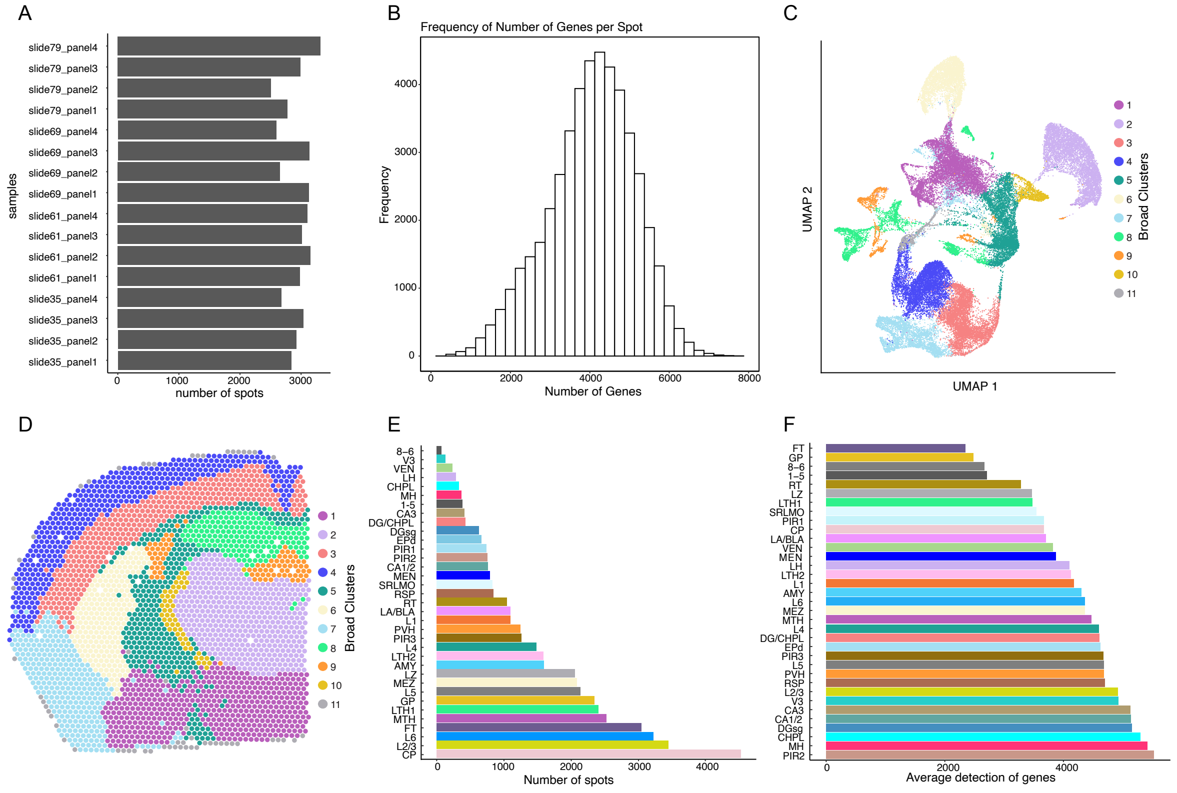

**Fig. S1. General QC measures and broad clustering.** (**A**) Total number of spots per slide. X-axis shows the number of spots and y-axis individual samples. Overall, the number of spots is relatively similar across samples. (**B**) Frequency of the number of genes per spot across all samples following a Gaussian distribution. X-axis depicts the number of genes, and y-axis represents the frequency. (**C**) Uniform Manifold Approximation and Projection for Dimension Reduction (UMAP) plot indicating the clusters based on PCA analysis and (**D**) 2D reconstruction of 11 distinct broad clusters. Colors indicate different clusters. (**E**) The total number of spots per brain region across all samples indicates that the CP is the relative largest brain region and thus receives more sequencing reads compared to other regions. (**F**) The average number of genes detected per brain region. Most brain regions share a similar number of genes.

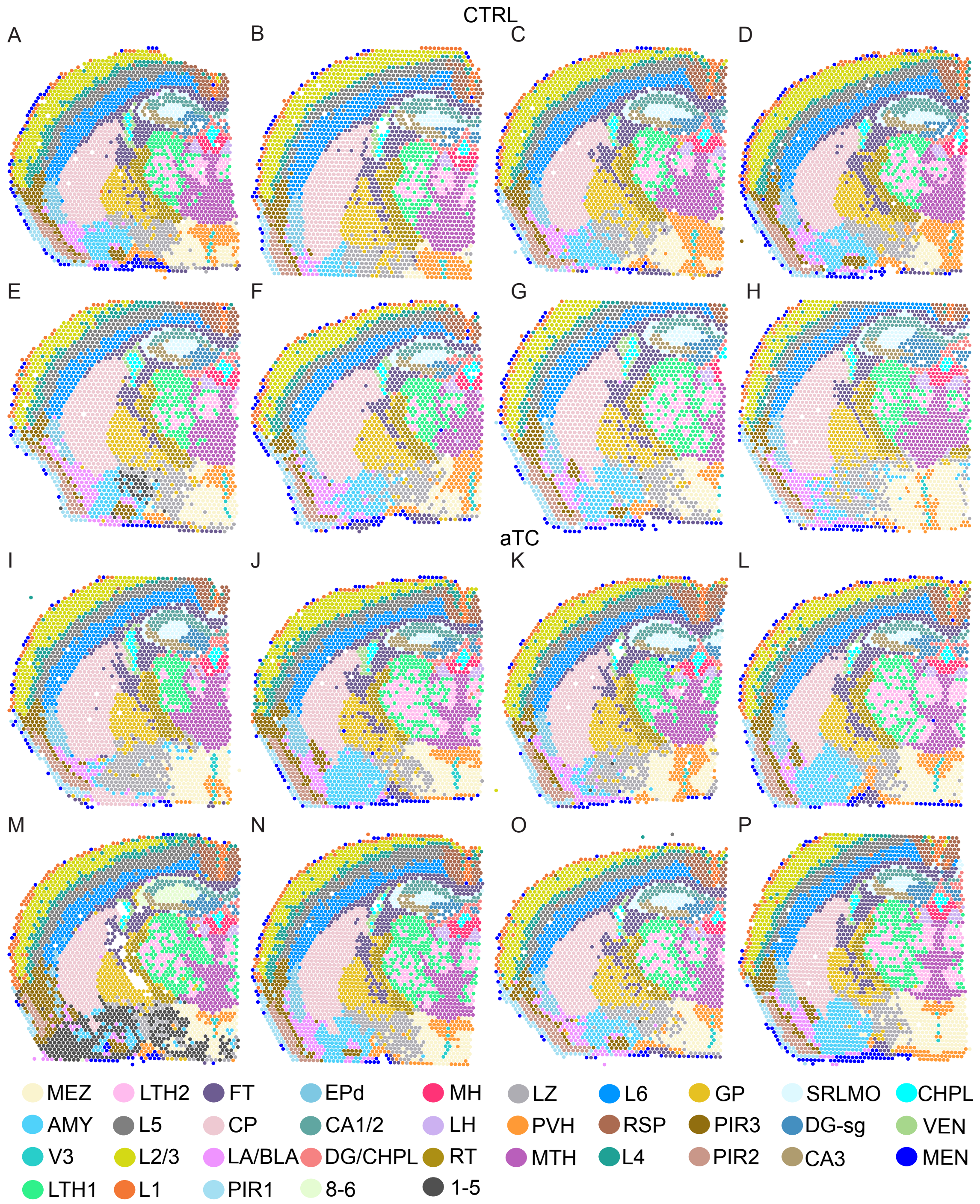


**Fig. S2. Sub-clustering resolves the mouse brain anatomy across all samples.** 2D reconstruction of the mouse brain anatomy using unsupervised clustering with the Leiden algorithm depicted on all samples. (**A-H**) Control samples and (**I-P**) aTC samples. Abbreviations for brain regions derived from ABA. MEZ: Hypothalamic medial zone, LZ: Hypothalamic lateral zone, AMY: Amygdala, PVH: Paraventricular hypothalamic nucleus, V3: third ventricle, MTH: Medial thalamus, LTH1: Lateral thalamus 1, LTH2: Lateral thalamus 2, L6: Cortical layer 6, L5: Cortical layer 5, RSP: Retrosplenial cortex, L2/3: Cortical layer L2/3, L4: Cortical layer 4, L1: Cortical layer 1, FT: Fiber tract, GP: Globus pallidus, CP: Caudoputamen, PIR3: Piriform area polymorph layer 3, LA/BLA: Lateral amygdalar nucleus and basolateral amygdalar nucleus, PIR2: Piriform area polymorph layer 2, PIR1: Piriform area polymorph layer 1, EPd: Endopiriform nucleus, SRLMO: includes stratum radiatum, stratum lacunosum-moleculare, stratum oriens and stratum lucidum, CA1/2: Hippocampal region CA1 and CA2, DGsg: Dentate gyrus granule cell layer, DG/CHPL: Dentate gyrus and choroid plexus, CA3: Hippocampal region CA3, MH: Medial habenula, CHPL: Choroid plexus, LH: Lateral habenula, VEN: Ventricle, RT: Reticular nucleus of the thalamus, MEN: Meninges, 1-5: Cluster 1-5 approximately located in the amygdala, 8-6: Cluster 8-6 approximately located in the hippocampus.

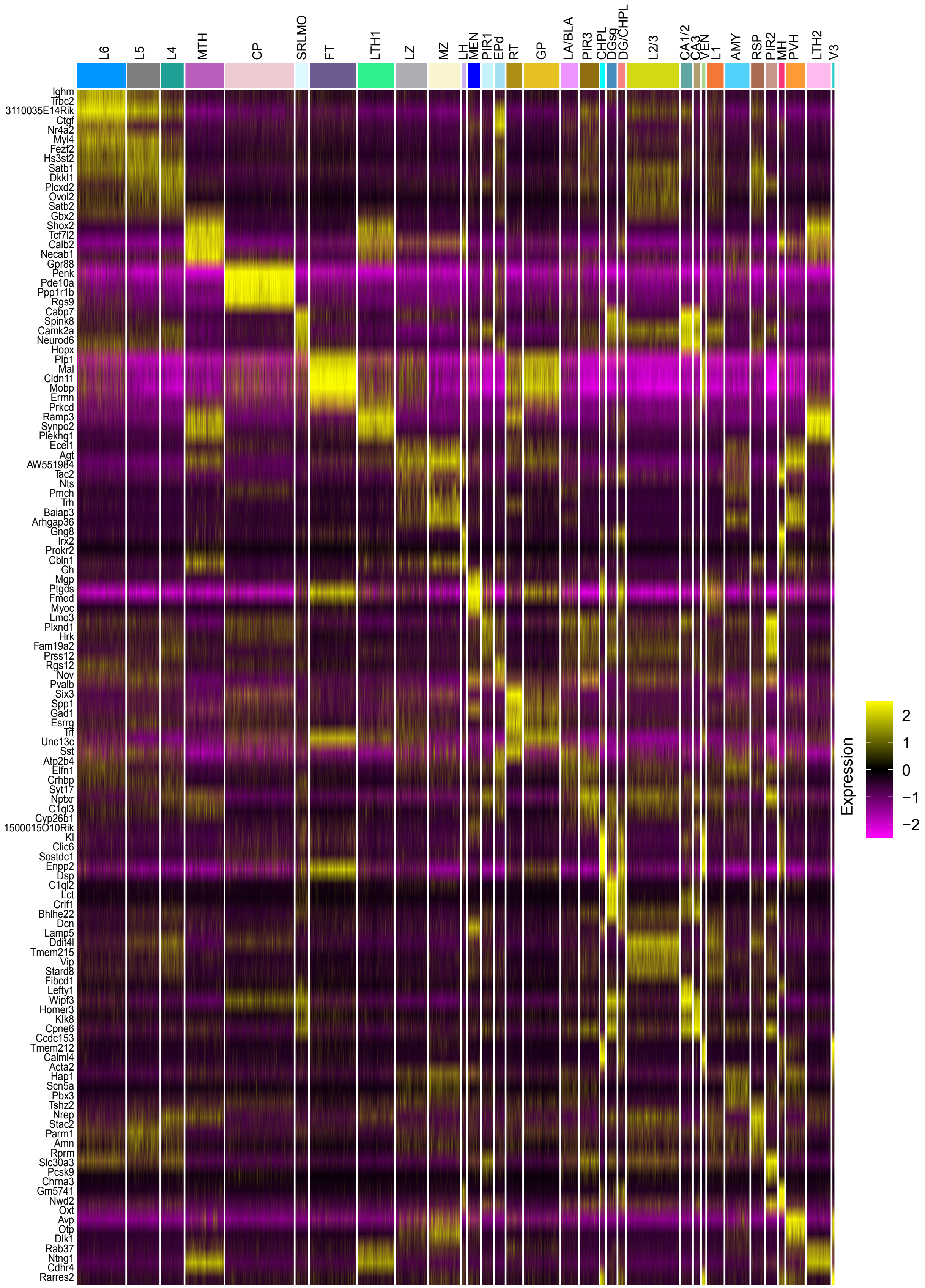

L6
L5
L4
MTH
CP
SRLMO
FT
LTH1
LZ
MZ
LH
MEN
PIR1
EPd
RT
GP
LA/BLA
PIR3
CHPL
DGsg
DG/CHPL
L2/3
CA1/2
CA3
VEN
L1
AMY
RSP
PIR2
MH
PVH
LTH2
V3
Ighm
Trbc2
3110035E14Rik
Ctgf
Nr4a2
Myl4
Fezf2
Hs3st2
Satb1
Dkkl1
Plcxd2
Ovol2
Satb2
Gbx2
Shox2
Tcf7l2
Calb2
Necab1
Gpr88
Penk
Pde10a
Ppp1r1b
Rgs9
Cabp7
Spink8
Camk2a
Neurod6
Hopx
Plp1
Mal
Cldn11
Mobp
Ermn
Prkcd
Ramp3
Synpo2
Plekhg1
Ecel1
Agt
AW551984
Tac2
Nts
Pmch
Trh
Baiap3
Arhgap36
Gng8
Irx2
Prokr2
Cbln1
Gh
Mgp
Ptgds
Fmod
Myoc
Lmo3
Plxnd1
Hrk
Fam19a2
Prss12
Rgs12
Nov
Pvalb
Six3
Spp1
Gad1
Esrrg
Ttr
Unc13c
Sst
Atp2b4
Elfn1
Crhbp
Syt17
Nptxr
C1ql3
Cyp26b1
1500015O10Rik
Kl
Clic6
Sostdc1
Enpp2
Dsp
C1ql2
Lct
Crlf1
Bhlhe22
Dcn
Lamp5
Ddit4l
Tmem215
Vip
Stard8
Fibcd1
Lefty1
Wipf3
Homer3
Klk8
Cpne6
Ccdc153
Tmem212
Calml4
Acta2
Hap1
Scn5a
Pbx3
Tshz2
Nrep
Stac2
Parm1
Amn
Rprm
Slc30a3
Pcsk9
Chrna3
Gm5741
Nwd2
Oxt
Avp
Otp
Dlk1
Rab37
Ntng1
Cdhr4
Rarres2
2
1
0
−1
−2
Expression

**Fig. S3. Heatmap of differentially expressed genes between brain regions.** Top 5 genes per region based on the Wilcoxon rank-sum test with $p_{FDR}<0.05$ are shown. X-axis ordered by brain region, y-axis shows top 5 genes per region. Yellow indicates upregulated gene expression ($log^2FC<0$) and in purple downregulated gene expression ($log^2FC>0$). Column width indicates the number of spots in a particular cluster.

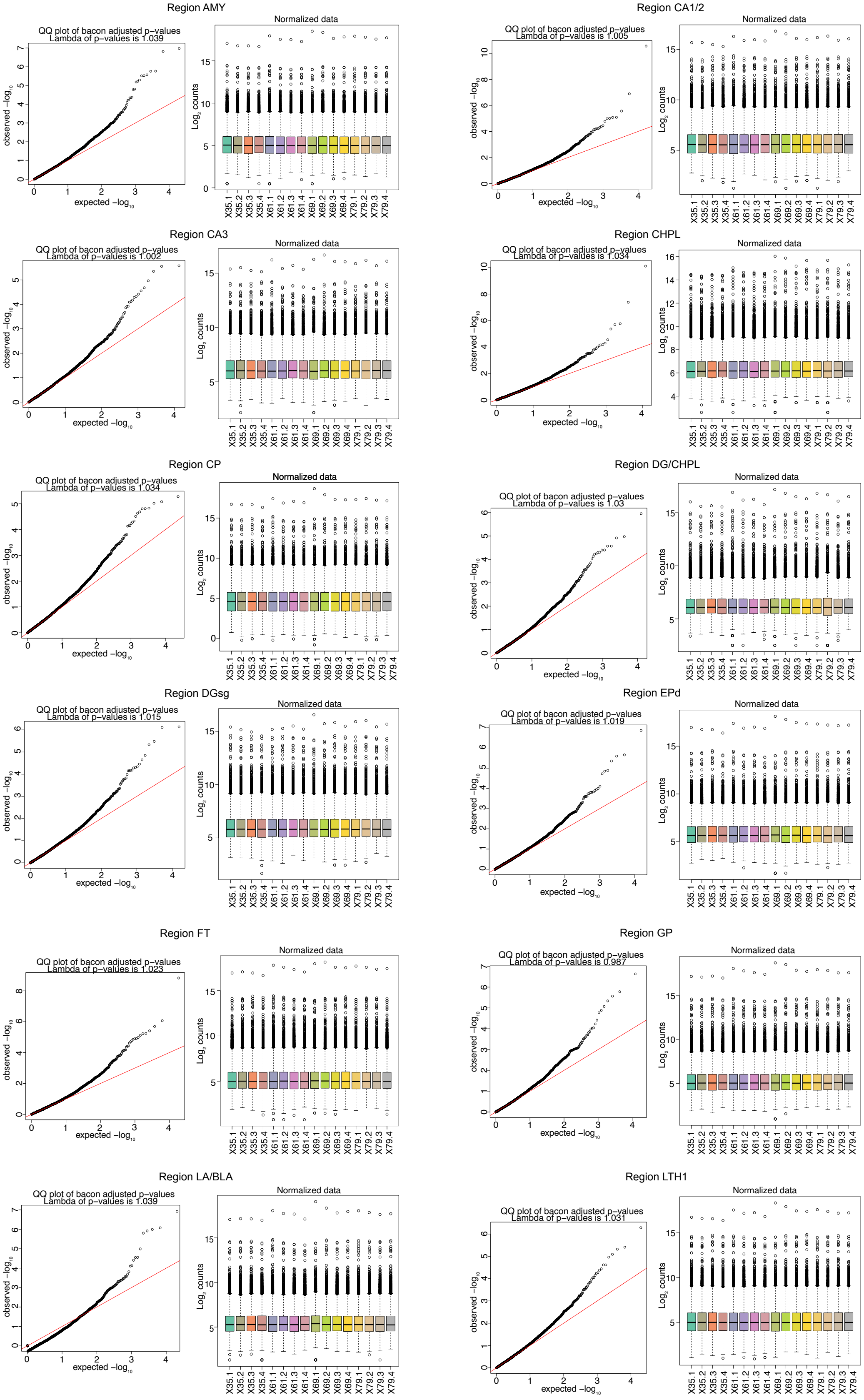
Region AMY
QQ plot of bacon adjusted p-values
Lambda of p-values is 1.039
Normalized data
Region CA1/2
QQ plot of bacon adjusted p-values
Lambda of p-values is 1.005
Normalized data
Region CA3
QQ plot of bacon adjusted p-values
Lambda of p-values is 1.002
Normalized data
Region CHPL
QQ plot of bacon adjusted p-values
Lambda of p-values is 1.034
Normalized data
Region CP
QQ plot of bacon adjusted p-values
Lambda of p-values is 1.034
Normalized data
Region DG/CHPL
QQ plot of bacon adjusted p-values
Lambda of p-values is 1.03
Normalized data
Region DGsg
QQ plot of bacon adjusted p-values
Lambda of p-values is 1.015
Normalized data
Region EPd
QQ plot of bacon adjusted p-values
Lambda of p-values is 1.019
Normalized data
Region FT
QQ plot of bacon adjusted p-values
Lambda of p-values is 1.023
Normalized data
Region GP
QQ plot of bacon adjusted p-values
Lambda of p-values is 0.987
Normalized data
Region LA/BLA
QQ plot of bacon adjusted p-values
Lambda of p-values is 1.039
Normalized data
Region LTH1
QQ plot of bacon adjusted p-values
Lambda of p-values is 1.031
Normalized data

Region LTH1

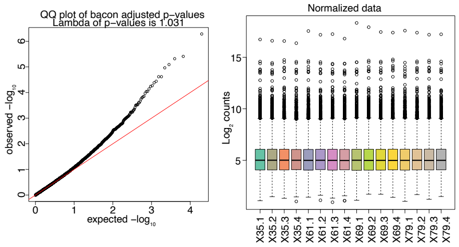


Region LTH2

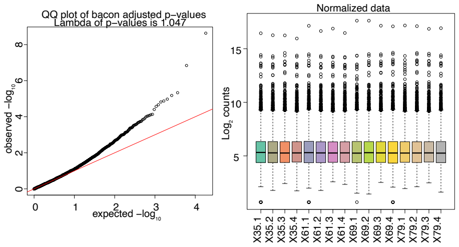


Region LZ

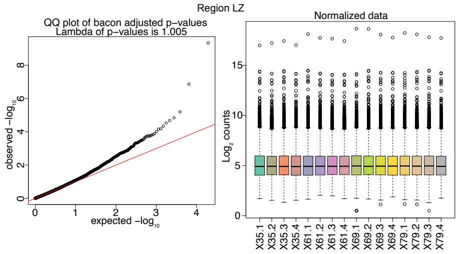


Region MEN

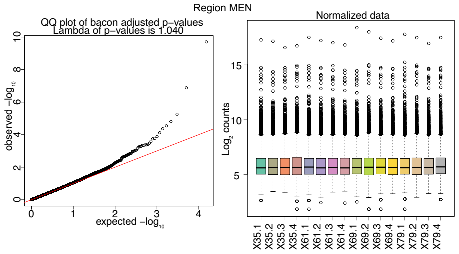


Region MEZ

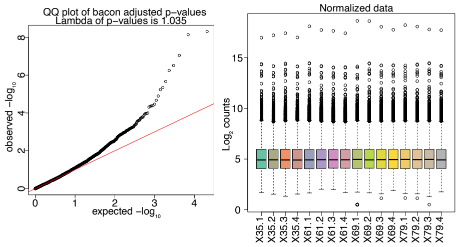


Region MH

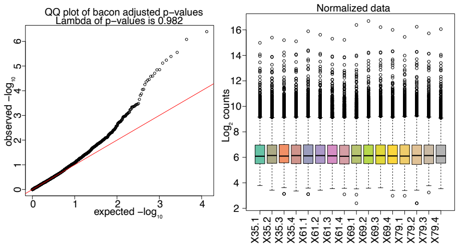


Region MTH

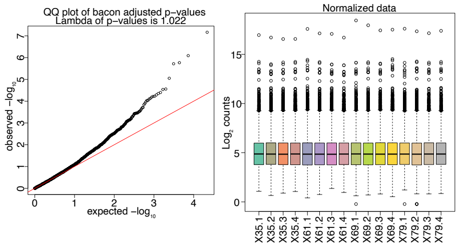


Region PIR1

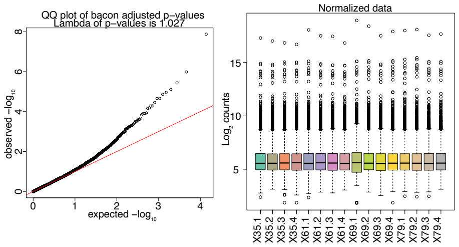


Region PIR2

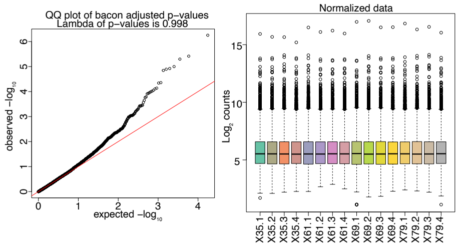


Region PIR3

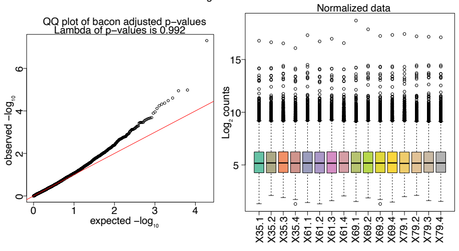


Region PVH

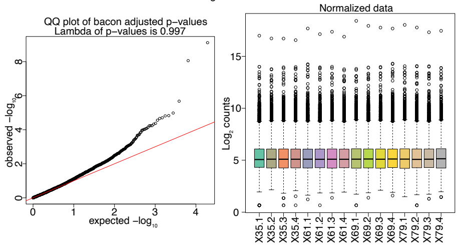


Region RSP

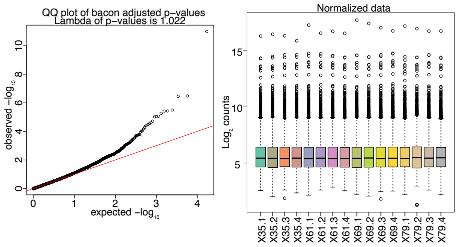

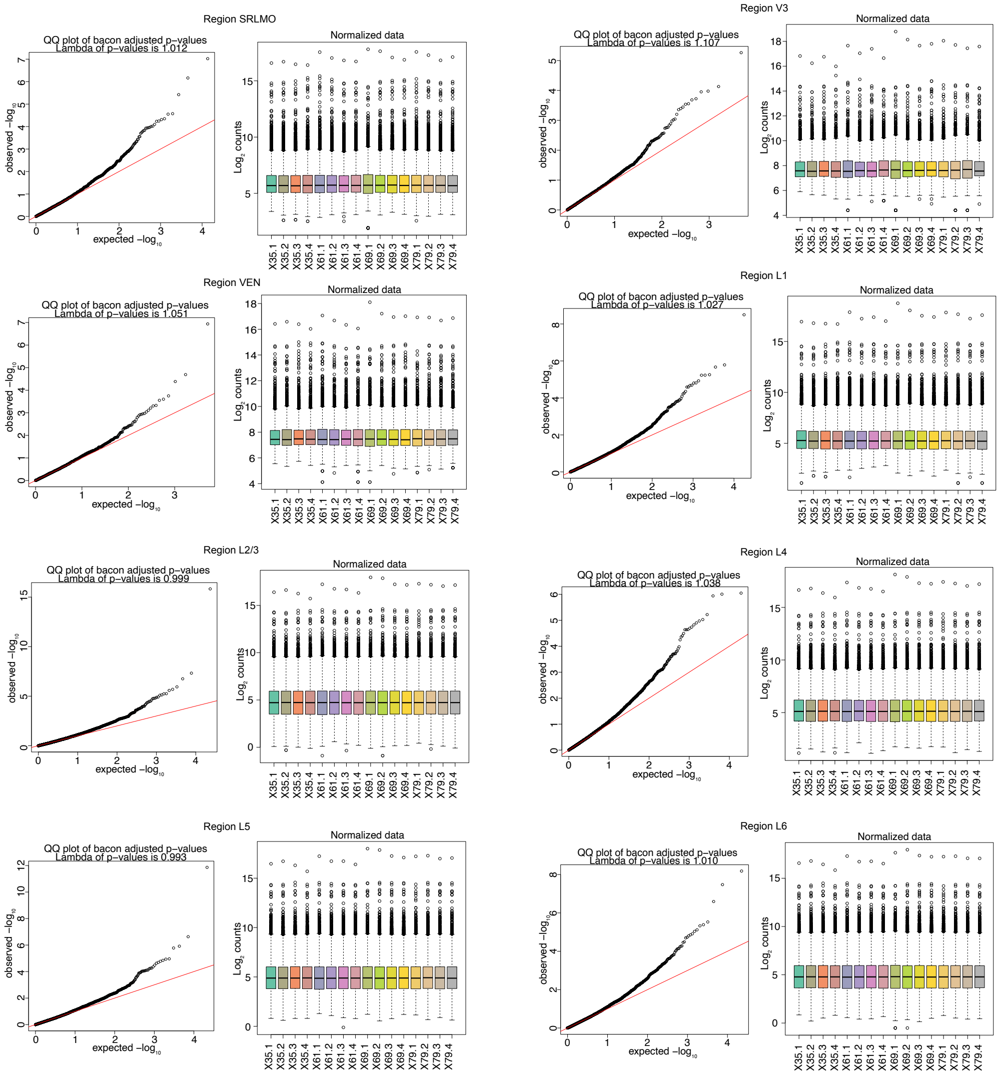


**Fig. S4. Quality control of DEG analysis.** Left, QQ plots that show the uniform distribution of the bacon-adjusted p-values per brain region. Lambda values indicate little to no inflation or deflation across analyses. The x-axis indicates the expected p-values and the y-axis the observed p-values. Right, normalized boxplots from voom-transformed data with the x-axis depicting the samples and, on y-axis, the normalized $log^2$ expression indicating correct normalization of the data across samples. Colors indicate different samples.

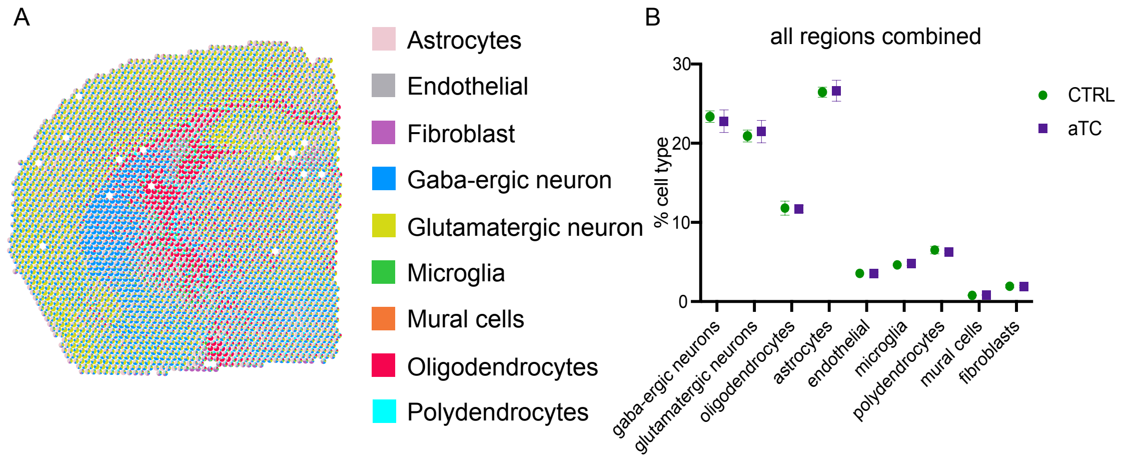


**Fig. S5. Deconvolution Visium spots.** (**A**) A representative sample deconvoluted with CARD, each spot represented by a pie-chart with the percentage of cell type contributing to that spot. Colors indicate the different cell types. (**B**) Boxplots of cell types across all samples. Green shows control and purple aTC samples (repeated measures two-way ANOVA, cell types x group interaction, $F_{8,\ 126}$=0.7869, p<0.05; Sidak's multiple comparisons posthoc test, p<0.05; n=8 per group). X-axis represents different cell types and the y-axis percentage of cell types across all samples, the data plotted is mean + SD.

A

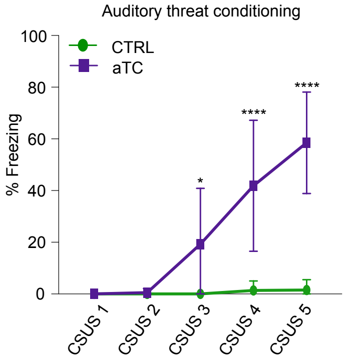


**Fig. S6. Auditory threat conditioning.** An independent cohort is subjected to auditory threat conditioning (n=7 CTRL, n=8 aTC). aTC data displaying the 5 tone-shock pairings (conditioned stimulus – unconditioned stimulus (CSUS) 1-5, x-axis) and percentage of time spent freezing on the y-axis. aTC results in a significant increase in freezing % in the threat-conditioned group after the second tone-shock pairing compared to the tone-control group. The data plotted is mean + SD, * = $p<0.05$; **** = $p<0.0001$.

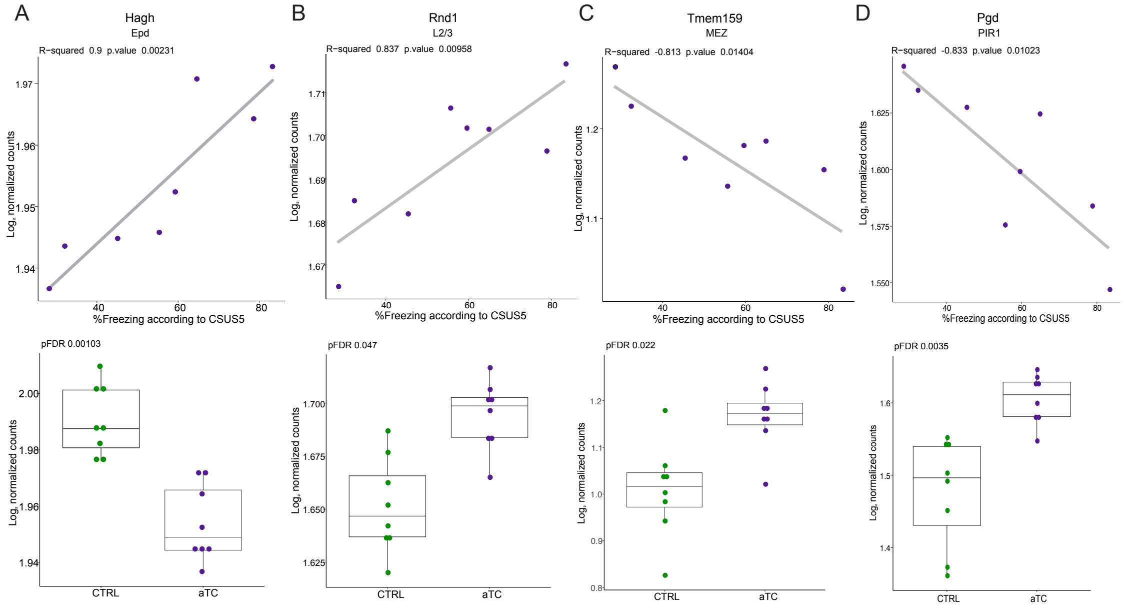


**Fig. S7. Correlation of DEGs with freezing behavior.** Example DEGs that correlate with freezing behavior (total 26/415). (**A-B**) *Hagh* and *Rnd1* correlate positively with freezing in the endopiriform nucleus or L2/3 respectively. (**C-D**) *Tmem159* and *Pgd* correlate negatively with freezing in the hypothalamic medial zone or piriform area polymorph layer 1 respectively. Pearson correlation: normalized counts ~ % freezing at CSUS 5, $p_{nominal}<0.05$.

**Supplemental Tables are provided as separate files.**

**Table S1. Region-specific marker genes.** The Wilcoxon-rank sum test was utilized to identify marker genes for each brain region. Each tab within the table corresponds to a different brain region. The average $log_2$ fold change was used to determine the marker genes for each specific region. The percentage of spots where a gene is detected within a particular cluster is denoted as pct.1 while pct.2 indicated the detection percentage of the gene in all other brain regions

**Table S2. Differentially expressed genes in response to auditory threat conditioning per brain region.** A pseudo-bulked linear model was applied to each brain region to compare control samples with auditory threat conditioning (aTC) samples. Each tab within the table corresponds to a different brain region. To control for the inflation or deflation of p-values, we used the bacon method, resulting in baconFDR values representing the adjusted $p_{FDR}$ values.

**Table S3. Correlation of gene expression with percentage freezing.** This table represents the correlation between gene expression levels and percentage freezing, measured at the $5^{th}$ tone-shock pairing. The correlation is denoted as an R-squared value, indicating the strength and direction of the relationship between gene expression and percentage freezing.

**Table S4. Enrichment analysis by brain region based on the differentially expressed genes.** Enrichment analysis was performed on differentially expressed genes in response to auditory threat conditioning (aTC) for each brain region, utilizing a pseudo-bulked linear model. Each tab represents a distinct brain region. The 'ID' and 'Description' columns denote the database source and the specific enriched term. ‘GeneRatio’ indicates the proportion of genes in the input list that overlap with the enriched term, while ‘BgRatio’ represents the ratio of genes in the input list relative to the total number of genes in the enrichment database.

**Table S5. Results from the weighted gene co-expression network analysis.** This table represents the results from the weighted gene co-expression network analysis (WGCNA). The tabs labeled “genes_darkgrey” and “genes_lightgreen” contain genes located in the darkgrey and lightgreen modules, respectively. The “enrichment_darkgrey” and “enrichment_lightgreen” tabs show the enrichment analysis results for genes within these modules. The “hubgenes_darkgrey” and “hubgenes_lightgreen” tabs indicate how well each gene is correlated within the module. The first column, labeled with “.r” represents the R-squared value; the gene with the highest R-squared value is identified as the hub gene of the respective module. The “.p” column indicates the p-value of the correlation.

**Table S6. Enrichment analysis of attenuated or over-patterned networks.** This table presents the enrichment analysis results for each attenuated or over-patterned brain region pair. Each tab contains the enrichment analysis outcomes specific to the brain-region pair.

**Table S7. Enrichment analysis of the over-patterned network of EPd-L4.** This table presents the enrichment analysis of the over-patterned network in the EPd-L4 regions. Genes are categorized based on their expression into either L4 or EPd brain region. A gene-set enrichment analysis was conducted, revealing the enrichment of SF-1 in cortical layer 4.
